# A Cell Penetrating Peptide from SOCS-1 Prevents Ocular Damage in Experimental Autoimmune Uveitis

**DOI:** 10.1101/315549

**Authors:** Chulbul M. Ahmed, Michael T. Massengill, Cristhian J. Ildefonso, Howard M. Johnson, Alfred S. Lewin

**Affiliations:** Department of Molecular Genetics and Microbiology, University of Florida Gainesville, FL 32610; Department of Microbiology and Cell Science, University of Florida Gainesville, FL 32610

**Author notes:** Corresponding author: Alfred S. Lewin, Ph.D., University of Florida, Department of Molecular Genetics and Microbiology, P.O. Box 100266, Gainesville, FL 32610.

**Keywords:** Immune suppression, autoimmune uveitis, retinal pigment epithelial cells, mouse model, ocular inflammation

## Abstract

We describe an immunosuppressive peptide corresponding to the kinase inhibitory region (KIR) of the intracellular checkpoint protein suppressor of cytokine signaling 1 (SOCS-1) that binds to the phospho-tyrosine containing regions of the tyrosine kinases JAK2 and TYK2 and the adaptor protein MAL, and thereby inhibits signaling downstream from these signaling mediators. The peptide, SOCS1-KIR, is thus capable of downregulating overactive JAK/STAT or NF-kB signaling in somatic cells, including those in many compartments of the eye. Attachment of poly-arginine to this peptide (R9-SOCS1-KIR) allows it to penetrate the plasma membrane in aqueous media. R9-SOCS1-KIR was tested in ARPE-19 cells and was found to attenuate mediators of inflammation by blocking the inflammatory effects of IFNγ, TNFα, or IL-17A. R9-SOCS1-KIR also protected against TNFα or IL-17A mediated damage to the barrier properties of ARPE-19 cells, as evidenced by immunostaining with the tight junction protein, zona occludin 1 (ZO-1), and measurement of transepithelial electrical resistance (TEER). Experimental autoimmune uveitis (EAU) was generated in B10.RIII mice using a peptide of interphotoreceptor retinal binding protein (IRBP^161-180^) as immunogen. Topical administration of R9-SOCS1-KIR protected ocular structure and function as seen by fundoscopy, optical coherence tomography (OCT), and electroretinography (ERG). The ability R9-SOCS1-KIR to suppress ocular inflammation and preserve barrier properties of retinal pigment epithelium makes it a potential candidate for aqueous treatment of autoimmune uveitis.

**Highlights:**

- peptide corresponding to the kinase inhibitory region of SOCS-1 linked to poly-arginine (R9-SOCS1-KIR) and its inactive control peptide were chemically synthesized.
- R9-SOCS1-KIR attenuated pro-inflammatory effects of IFNγ, TNFα, and IL-17 in ARPE-19 cells, thus showing a simultaneous inhibition Th1 and Th17 cell functions.
- Damage to barrier properties of ARPE-19 cells caused by TNFα or IL-17 was prevented in the presence of R9-SOCS1-KIR.
- Topical administration of R9-SOCS1-KIR prevented ocular damage in a mouse model of experimental autoimmune uveitis.

## 1. Introduction

Uveitis is a complex group of sight threatening diseases that can result from infection, systemic inflammation, or an autoimmune response. Uveitis is estimated to cause 10-15% of all cases of blindness in the US (Miserocchi et al., 2013). In recent years, host-microbiota interaction has been shown to contribute to both infectious and non-infectious uveitis, suggesting a common source of the two types (Heissigerova et al., 2016). Autoimmune uveitis activates both the innate and adaptive immune responses, where the causative ocular antigens are well characterized, and the multifactorial nature of the disease has been examined (reviewed in (Perez and Caspi, 2015)). Autoimmune uveitis can be restricted to the eye (examples include idiopathic uveitis, sympathetic ophthalmia, or birdshot retinochoroidopathy), or it can be a systemic response (e.g., sarcoidosis, Behcet’s disease, Vogt-Koyanagi-Harada syndrome). Experimental autoimmune uveitis (EAU) is induced in experimental animals by immunizing with uveitogenic ocular antigens. These animal models have been invaluable tools in understanding the mechanism of disease as well as in developing therapies (Caspi, 2010). It is well established that either the Th1 or the Th17 response is capable of causing autoimmune uveitis. Neutralization of Th1 response leads to an elevated Th17 response, and a deficiency of Th17 response leads to an elevated Th1 response (Luger et al., 2008). Thus, a simultaneous blockage of both Th1 and Th17 responses is crucial to the treatment of autoimmunity.

The current treatments for non-infectious uveitis include corticosteroids, general immunosuppressants, or specific antibodies. Corticosteroid treatment is given topically, through intraocular injection, or systematically. Although initial inflammation is suppressed, continued treatment with corticosteroids is associated with development of cataracts, glaucoma, retinopathy, and activation of herpes simplex virus. Immunosuppressant agents, such as cyclosporine A (CsA), a T cell targeting drug that blocks IL-2 signaling has been employed to combat ocular inflammation. FK-506 (tacrolimus) and rapamycin, which also target IL-2 signaling, have also been used. This line of treatment is discouraged, because of the involvement of IL-2 in maintenance of Tregs, since decreased levels of Tregs are undesirable. Furthermore, these treatments are also not advised for a prolonged periods. Antibodies targeting tumor necrosis factor α (TNFα) or IL-1 that can suppress inflammatory signaling pathways are currently being tested (Schwartzman, 2016). Therefore, effective treatments for uveitis is an unmet need.

The extent and duration of an immune response is negatively regulated by a set of intracellular proteins called as <u>s</u>uppressors <u>o</u>f <u>c</u>ytokine <u>s</u>ignaling (SOCS) (Yoshimura et al., 2012). Amongst the known 8 SOCS proteins, SOCS1 and SOCS3 are particularly relevant to nearly all the cells where JAK/STAT and NF-kB signaling is utilized. SOCS1 contains a kinase inhibitory region (KIR), which spans the residues 53 to 68 of the SOCS1 protein. We have previously shown that the peptide SOCS1-KIR binds to the kinase activation loop containing phosphotyrosine residues of such important tyrosine kinases, as JAK2 and TYK2, and the toll-like receptor adaptor protein MAL (Jager et al., 2011). Owing to this property, the SOCS1-KIR peptide is capable of suppressing signaling downstream of these molecules. Attachment of palmitoyl-lysine allowed this peptide (lipo-SOCS1-KIR) to gain access across the plasma membrane. We showed that this peptide was capable of protecting against experimental autoimmune encephalomyelitis (EAE), a mouse model of multiple sclerosis (Jager et al., 2011). Here, we report the synthesis and use of a water soluble version of SOCS1-KIR, denoted as R9-SOCS1-KIR. R9 represents the 9 arginines attached to SCOS1-KIR that causes both water solubility and cell penetration. R9-SOCS1-KIR is likely to be less immunogenic than lipo-SOCS1-KIR because it lacks the lipid moiety. In this report, we describe the ability of R9-SOCS1-KIR to suppress inflammatory responses emanating from IFNγ, TNFα, or IL-17, and protection against the loss of transepithelial electrical resistance (TEER) in a human retinal pigment epithelial cell line, ARPE-19. Since multiple sclerosis and autoimmune uveitis share several common features and can in some cases be overlapping (Gordon and Goldstein, 2014; Messenger et al., 2015), we tested R9-SOCS1-KIR in a mouse model of experimental autoimmune uveitis (EAU), and found it to be capable of protecting against the ocular damage in EAU by simple application as eye drops.

## 2. Materials and Methods

### 2.1. Cell culture

ARPE-19 cells (ATCC, 2302) were grown in DMEM/F-12 medium containing 10% FBS and 1% each of Penicillin, Streptomycin in a humidified incubator at 37 °C. THP-1 cells (ATCC, TIB-202) were grown in RPMI medium with 10% FBS and 1% each of Penicillin, Streptomycin.

### 2.2. Peptide synthesis

Conventional fluorenylmethyl-oxycarbonyl (FOMC) chemistry was used for the synthesis of peptides on an Applied Biosystems 431A automated peptide synthesizer, as described previously (Szente et al., 1994). R9-SOCS1-KIR had the following sequence: RRRRRRRRRDTH**F**RT**F**RSHSDYRRI. The critical phenylalanines shown in bold were replaced by alanines in an inactive control peptide, R9-SOCS1-KIR2A, which had the following sequence: RRRRRRRRRDTH**A**RT**A**RSHSDYRRI. Peptides were characterized by mass spectrometry and purified by high performance liquid chromatography (HPLC). Peptides were reconstituteded in PBS under sterile conditions before use.

### 2.2. Immunohistochemistry

ARPE-19 cells were grown in 8 well chambered slides. After overnight growth in regular DMEM/F12 medium with 10% FBS, the medium was changed to DMEM/F12 medium with 1% FBS. While testing for the effects of IFNγ and IL-17A, cells were pre-treated with R9-SOCS1-KIR or the control peptide R9-SOCS1-KIR2A at 20 μM for 3 hrs followed by treatment with IFNγ (1 ng/ml), or IL-17A (50 ng/ml) for 30 min. In experiments where ARPE-19 cells were differentiated to cuboidal morphology, cells were grown in 1% FBS containing medium with a change of media twice per week for 4 weeks. Cells were pre-treated with R9-SOCS1-KIR or the control peptide R9-SOCS1-KIR2A at 20 μM for 3 hrs followed by treatment with TNFα (10 ng/ml), or IL-17A (50 ng/ml) for 48 hr. Cells were fixed with 4% paraformaldehyde for 30 min at room temperature and washed with PBS, followed by permeabilization using 1% Triton X-100 in PBS for 30 min at room temperature. Cells were then blocked in 10% normal goat serum in PBS containing 0.5% Triton X-100 for 30 min at room temperature followed by washing in 0.2% Triton X-100 in PBS (wash buffer). Rabbit polyclonal antibody to pSTAT1α (Santa Cruz Biotechnology), pSTAT3 (Cell Signaling), p65 (Cell Signaling), or ZO-1 (Invitrogen) (1:200 dilution) were added and incubated overnight at 4° C, followed by washing. Cy-3 conjugated anti-rabbit secondary antibody (Invitrogen, 1:300 dilution), or Alexa Fluor 488 conjugated secondary antibody (Invitrogen, 1:300 dilution) were added and incubated for 30 min at room temperature, followed by washing. After the addition of mounting media, cells were covered with a cover slip and imaged under a Keyence BZ-X700 fluorescence microscope.

### 2.3. RNA extraction and qPCR

ARPE-19 cells were seeded in a 12 well plate at 70% confluence. The next day, the medium was changed to DMEM/F-12 containing 1% FBS. Cells were pre-treated with 20 μM R9-SOCS1-KIR for 3 hr, followed by treatment with 10 ng/ml of TNFα for 1 hr. Cells were then washed with PBS and total RNA was extracted using the RNeasy Mini Kit from QIAGEN, following the manufacturer’s instructions. One microgram of RNA was used to synthesize first strand cDNA using the iScript cDNA synthesis kit from Bio-Rad (Hercules, CA). The following PCR primers, synthesized by IDT (Coralville, Iowa) were used. IL-1β (For: CTCGCCAGTGAAATGATGGCT, Rev: GTCGGAGATTCGTAGCTGGAT), IL-6 (For: CTTCTCCACAAGCGCCTTC, Rev: CAGGCAACACCAGGAGCA), CCL-2 (For: CTCATAGCAGCCACCTTCATTC, Rev: TCACAGCTTCTTTGGGACACTT), and β-actin (For: AGCGAGCATCCCCCAAAGTT, Rev: GGGCACGAAGGCTCATCATT). The PCR reaction mixture contained cDNA template, SsoFastEvaGreen Super mix containing SYBR green (Bio-Rad (Hercules, CA), and 3 μM gene specific primers. After denaturation at 95° C for 2 min, 40 cycles of reaction including denaturation at 94° C for 15 sec followed by annealing at 60° C for 30 sec were carried out using C1000 thermal cycler CFX96 real-time system (Bio-Rad). Gene expression was normalized to beta actin. Relative gene expression was compared with untreated samples and determined using the CFX96 software from Bio-Rad.

### 2.4. NF-κB promoter activity

A plasmid, pNF-κB-Luc that contains NF-κB promoter linked to firefly luciferase and another plasmid with a constitutively expressing thymidine kinase promoter driven Renilla luciferase (pRL-TK-Luc) were obtained from Promega. pRL-TK-Luc served as an internal control for the efficiency of transfection. ARPE-19 cells were seeded in 12 well plates at 90% confluency. The next day, cells were taken in a serum free media and treated with SOCS1-KIR or SOCS1-KIR2A at 20 μM for 3 hr followed by treatment with TNFα (10 ng.ml) for 4 hr. Cells were transfected by using 2 μg of pNF-kB-Luc, 10 ng of pRL-TK and 2 μg of NovaFector reagent (VennNova) per well for 24 hrs. Cell lysates were used to measure firefly and Renilla luciferase with a dual luciferase assay kit (Promega). Relative luciferase units were calculated by dividing firefly luciferase activity by Renilla luciferase activity in each sample.

### 2.5. Measurement of transepithelial electrical resistance

ARPE19 cells were plated on 24-well transwell inserts (Greiner-Bio-one, surface area 33.6 mm^2^, 0.4 μm pore size) in DMEM/F-12 with 1% FBS for 4 weeks, with a change of media twice per week. Cells were pre-treated with R9-SOCS1-KIR at 20 μM for 3 hr followed by treatment with TNFα (10 ng/ml), or IL-17A (50 ng/ml) for 48 hr. Transepithelial electrical resistance was measured using an EVOM2 voltohmmeter (World Precision Instruments), according to the manufacturer’s instructions. The plates containing the transwells were allowed to equilibrate to room temperature. The inserts were removed one at a time and placed into the EVOM2 chamber filled with DMEM/F-12 basal media. Net TEER measurements were calculated by subtracting the value of a blank transwell filter from the value of each filter with plated cells.

### 2.6. Induction of experimental autoimmune uveitis

B10.RIII mice (Jackson Lab) were housed in standard caging in a room with a 12h light: 12 hour dark cycle. All animal procedures were approved by the University of Florida Institutional Animal Care and Use Committee and were conducted in accordance with the Association for Research in Vision and Ophthalmology (ARVO) Statement for the Use of Animals in Ophthalmic and Vision Research. B10.RIII mice (female, 6 to 8 weeks old, n = 8) were immunized with 100 μg human IRBP160-181 (GenScript) in 100 μl PBS emulsified in 100 μl of complete Freund’s adjuvant administered at the base of the tail and s.c. on the thighs. Administration of R9-SOCS1-KIR dissolved in PBS (15 μg in 2 μl per eye), or PBS (2 μl per eye) began two days before immunization and continued daily for three weeks after immunization.

### 2.7. Fundoscopy and Spectral domain optical coherence tomography (SD-OCT)

For certain procedures, such as fundoscopy and spectral domain optical coherence tomography (SD-OCT), mice were anesthetized with a mixture of ketamine (95mg/kg) and xylazine (5-10 mg/kg) by intraperitoneal injection. Local anesthesia for the cornea was provided by 1 drop of proparacaine (1%). Fundus images were taken using a Micron III retinal imaging microscope (Phoenix Research Laboratories, Pleasanton, CA, USA). Spectral domain optical coherence tomography (SD-OCT) employed an Envisu ultra high resolution instrument (Bioptigen, Research Triangle Park, NC) at monthly intervals. Mice were dilated and anesthetized as described above for ERG analysis. We collected 250 linear B-scans and 25 images were averaged to minimize the background noise and to achieve a better resolution. Four measurements equidistant from the optical nerve head were recorded from each eye and were analyzed to determine the difference in ONL thickness. The results from each eye were averaged and the means were compared between time points.

Infiltrating cells in the vitreous were quantitated by using Image J software from NIH (URL:https://imagej.nih.gov/ij/). From the OCT image, the area representing the vitreous was delineated. The spots therein representing the infiltrating cells were converted into binary images and the software generated the total number of cells in the selected area. The number of cells in three different layers (B-scans) of the stack were used to average the number of particles present in the field of view. The number of cells in both eyes were counted from these digital images and averaged. A comparison between the cells treated with PBS or SOCS1-KIR was made.

### 2.8 Electroretinography (ERG)

Mice were dark adapted overnight and anesthetize as above. In dim red light a 2.5% phenylephrine solution was applied. Gold wire loop contact lenses were placed on each cornea using a drop of CVS brand lubricant solution to maintain corneal hydration. A ground electrode was placed subcutaneously in a hind leg, and a silver wire reference electrode was placed subcutaneously between the eyes. Responses were recorded from both eyes using an LKC UTAS Visual Electrodiagnostic System with a BigShot^TM^ full-field dome (LKC, Gaithersburg, MD). Scotopic ERGs were elicited with 10 msec flashes of white light at 0 dB (2.68 cds/m^2^), −10 dB (0.18 cds/m^2^), −20 dB (0.02 cds/m^2^) with appropriate delay between flashes. Five to 10 scans were averaged at each light intensity. This was followed by a two minute white light bleach in room light then a white flash at 1.0 cd sec/m^2^ intensity was used to elicit a cone response. The a-wave amplitudes were measured from baseline to the peak in the cornea-negative direction, and b-wave amplitudes were measured from cornea-negative peak to major cornea-positive peak. Both a-wave and b-wave amplitudes were measured before immunization with IRBP peptide and 4 weeks after immunization.

## 3. Results

### 3.1. R9-SOCS1-KIR prevents IFNγ, IL-17A, and TNFα-mediated activation and nuclear translocation of STAT1α, STAT3, and NF-κBp65

The retinal pigment epithelial (RPE) cells play a crucial role in maintaining the blood-ocular barrier (Fuhrmann et al., 2014; Simo et al., 2010). A compromise in the RPE structure results in infiltration of immune cells into the eye and reduced phagocytosis of the photoreceptor outer segment from the retina. We have used a spontaneously derived human RPE cell line, ARPE-19 cells, as a model system to test the prevention of inflammatory responses in cell culture by R9-SOCS1-KIR. Since IFNγ, IL-17A and TNFα are the major disease promoters of autoimmune uveitis (Luger et al., 2008), we tested their effect on ARPE-19 cells grown in 8 well microscopic chamber slides. IFNγ binding to its receptor results in phosphorylation of JAK2 (pJAK2). Phosphorylated JAK2 along with the receptor ligand complex recruits STAT1α, which upon phosphorylation (pSTAT1α) is translocated to the nucleus to activate target genes. Since SOCS1-KIR peptide inhibits phosphorylation of JAK2 (Jager et al., 2011; Waiboci et al., 2007), downstream signaling from this is prevented. Treatment of cells with IFNγ (1 ng/ml) for 30 min resulted in increased nuclear translocation of pSTAT1α, as seen by staining with an antibody to pSTAT1α and fluorescence microscopy (**Figure 1A**). Cells that were pre-treated with R9-SOCS1-KIR at 20 μM for 3 hrs followed by IFNγ (1 ng/ml for 30 min) treatment, or untreated cells, failed to show the nuclear translocation of pSTAT1α. Pre-treatment with the inactive control peptide R9-SOCS1-KIR2A at 20 μM for 3 hrs followed by treatment with IFNγ was not able to prevent the nuclear translocation of pSTAT1α, suggesting the specificity of action of R9-SOCS1-KIR. Thus, treatment with R9-SOCS1-KIR was capable of preventing IFNγ signaling in ARPE-19 cells.

**Figure 1.**
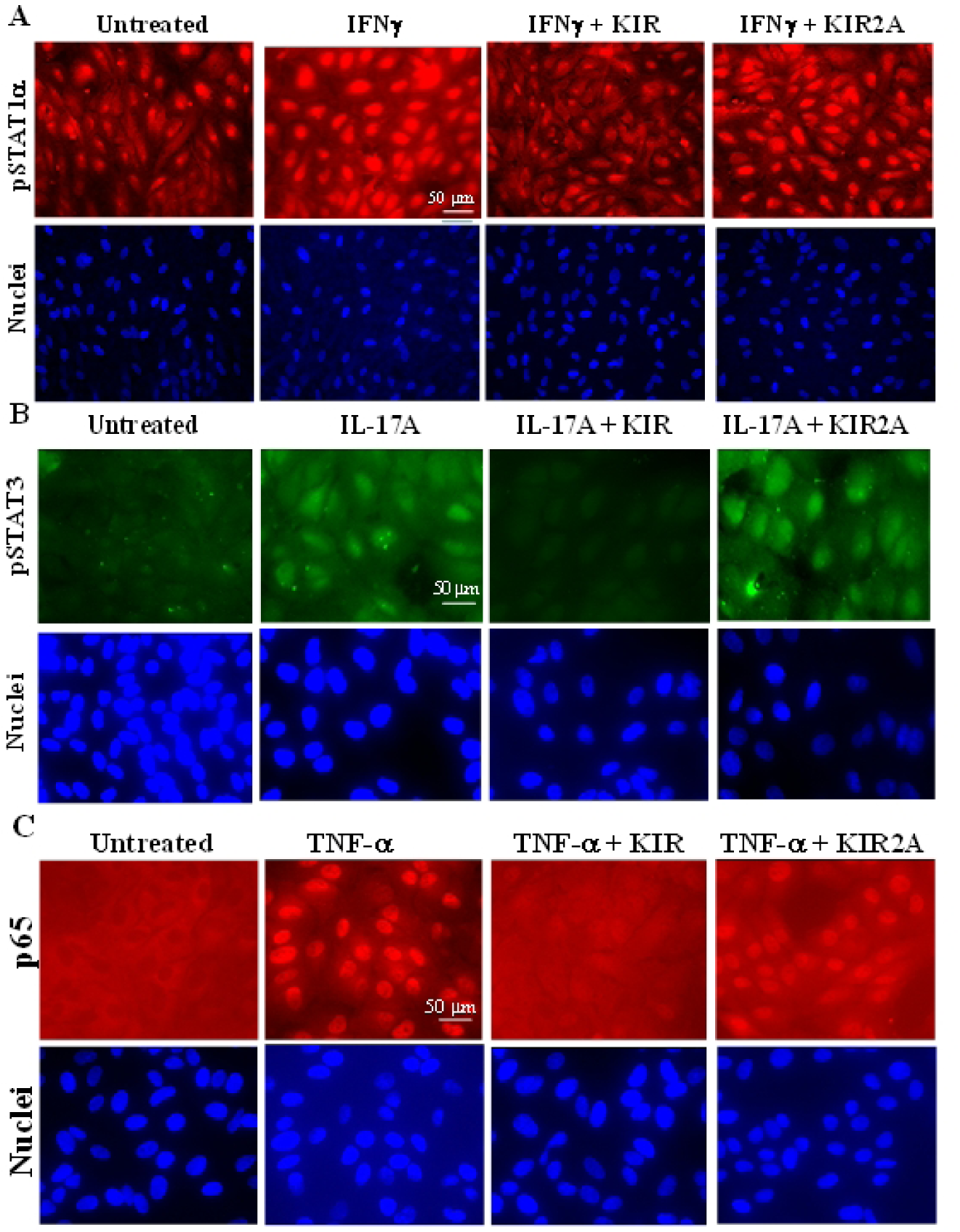
R9-SOCS1-KIR prevents nuclear translocation of activated STAT1α**(A)**, STAT3 (**B**), and NF-κB p65 subunit (**C**). ARPE-19 cells were seeded in serum free media in an 8 well cell culture slide and grown overnight. ARPE-19 cells were seeded in 8 well cell culture slide and grown overnight. Cells were taken in serum free media and pre-treated with R9-SOCS1-KIR (KIR) peptide, or its inactive control peptide R9-SOCS-KIR2A (KIR2A) at 20 μM for 3hr, followed by treatment for 30 min each with IFNγ (1 ng/ml) (**A**), IL-17A (50 ng/ml)(**B**), or TNFα (10 ng/ml) (**C**). Untreated cells or those treated with the cytokine alone were used alongside. Cells were stained with an antibody to pSTAT1α (**A**), pSTAT3 (**B**), or NF-kB p65 subunit (**C**), followed by staining with fluorescence conjugated secondary antibody and imaged in a fluorescence microscope.

Since IL-17 is another inflammatory cytokine involved in autoimmune diseases, including uveitis (Horai and Caspi, 2011), we tested the effect of R9-SOCS1-KIR on IL-17A signaling. IL-17A has been shown to act through the tyrosine kinase JAK2 which together with the ligand receptor complex phosphorylates STAT3, causing its translocation into the nucleus (Xu et al., 2014). ARPE-19 cells were treated with R9-SOCS1-KIR or R9-SOCS1-KIR2A at 20 μM for 3 hr followed by addition of IL-17A at 50 ng/ml for 30 min. Cells were then stained with an antibody to pSTAT3 and DAPI (for nuclei) and imaged in a fluorescence microscope (**Figure 1B**). IL17-A treatment resulted in the nuclear translocation of pSTAT3 which was prevented by R9-SOCS1-KIR, while the control peptide (KIR2A) had no effect on the nuclear translocation of pSTAT3. Thus, R9-SOCS1-KIR can block the effects of both IFNγ and IL1-7.

TNFα activates the transcription factor NF-κB, which culminates in the nuclear translocation of p65, the active subunit of NF-κB. This activation was followed by fluorescence microscopy. Addition of TNFα to ARPE-19 cells at 10 ng/ml for 30 min resulted in nuclear translocation of p65, which was suppressed when cells were pretreated with R9-SOCS1-KIR, and not by the inactive control peptide R9-SOCS1-KIR2A (**Figure 1C**), which indicates that R9-SOCS1-KIR can downregulate NF-κB promoter activity.

### 3.2. SOCS1-KIR attenuates inflammatory injury caused by TNFα in ARPE-19 cells

Tumor necrosis factor α (TNFα) is also associated with the onset of uveitis (Al-Gayyar and Elsherbiny, 2013). We thus investigated the effect of TNFα on the ARPE-19 cells and determined if prior treatment with the SOCS1-KIR would suppress the indicators of inflammation (**Table 1**). RNA from ARPE-19 cells treated as indicated was isolated and used for cDNA synthesis, followed by qPCR to quantitate RNA expression in target genes. The cytokine IL-1β was induced 190-fold at the RNA level as assayed by qPCR. Pre-treatment with R9-SOCS1-KIR resulted in a greater than 25% decrease in the level of IL-1β. Similarly, the chemokine CCL-2 and cytokine IL-6 were induced 44- and 6-fold, respectively, and this induction was reduced significantly (48% and 56%, respectively) by pre-treatment with R9-SOCS1-KIR. These cytokines and chemokine are associated with uveitis-related inflammation, recruitment of T cells and monocytes, and loss of barrier function of RPE. To verify the induction of IL1-β caused by treatment of ARPE-19 cells with TNFα and its suppression by R9-SOCS1-KIR, we carried out an ELISA assay (**Figure 2**). In response to TNFα, up to 80 pg/ml of IL-1β was secreted, and this level was reduced 4-fold by prior treatment with R9-SOCS1-KIR. A control peptide, SOCS1-KIR2A had no significant effect on the secretion of IL-1β in response to TNFα. The induction of IL-1β was also tested in a human monocytic cell line, THP-1 (**Figure 3**). THP-1 cells were treated with IFNγ (1 ng/ml) followed by treatment with *E. coli* lipopolysaccharide (LPS) (1 μg/ml) overnight. This combined treatment is needed to induce a pronounced response in THP-1 cells. About 55 pg/ml of IL-1β was secreted by treatment with IFNγ and LPS, and this level was reduced by one-half by the pre-treatment with R9-SOCS1-KIR, while the control peptide had an insignificant inhibitory effect. These results are consistent with the suppression nuclear localization of STAT1 by R9-SOCS1-KIR (**Figure 1**), and also points to the ability of R9-SOCS1-KIR to attenuate signaling from TLR4, since LPS acts through this receptor.

**Table 1.** TNFα mediated inflammatory responses are suppressed by SOCS1-KIR-R9. ARPE-19 cells at 90% confluency were seeded and grown overnight. They were changed into serum free media and treated with 20 μM SOCS1-KIR-R9 for 3 hrs, followed by treatment with TNFα (10 ng/ml) for 4 hrs. RNA was extracted from these cells and was used for cDNA synthesis followed by qPCR with primers for target genes indicated. Samples were run in triplicate and the relative expression versus β-actin was determined by qPCR. Values represent average fold change over the untreated samples ± s.d. Statistical significance for the three target genes compared between untreated, TNFα, or TNFα + SOCS1-KIR treated samples was < 0.0001, as measured by one way ANOVA followed by Tukey’s test.

| Target gene | Relative expression (compared with untreated) ± s.d. | | % Inhibition | P value |
| --- | --- | --- | --- | --- |
| | TNFα | TNFα + SOCS1-KIR | | |
| IL-1β | 1963 ± 28 | 144.0 ± 6.3 | 27 | 0.03 |
| CCL-2 | 44.3 ± 4.6 | 23.1 ± 2 | 48 | 0.013 |
| IL-6 | 5.9 ± 0.8 | 2.6 ± 0.28 | 56 | 0.001 |

**Figure 2.**
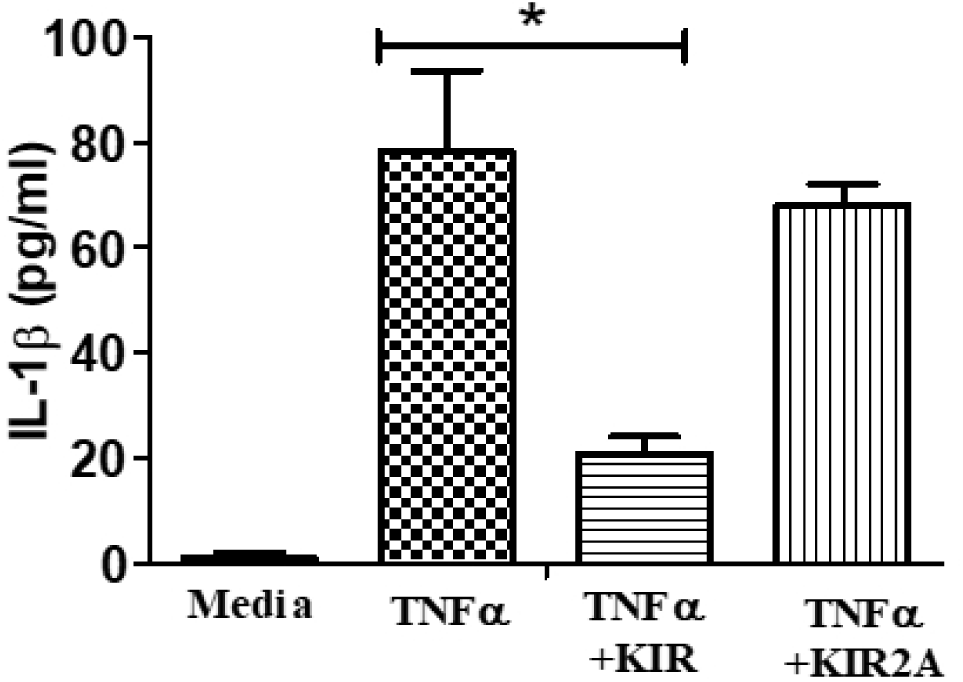
R9-SOCS1-KIR peptide suppresses the secretion of IL-1β from cells induced with TNFα. ARPE 19 cells were seeded in low serum medium in 12 well dishes and grown overnight. They were then treated with R9-SOCS1-KIR (KIR), or its control R9-SOCS1-KIR2A (KIR2A) peptide (20 μM) for 3 hrs. TNFα was added at 10 ng/ml and cells were grown for 18 hrs. Supernatants were harvested and used in triplicate for quantitation of IL-1β in an ELISA format. Bars represent the average of triplicates +/- s.d. *, p = 0.004.

**Figure 3.**
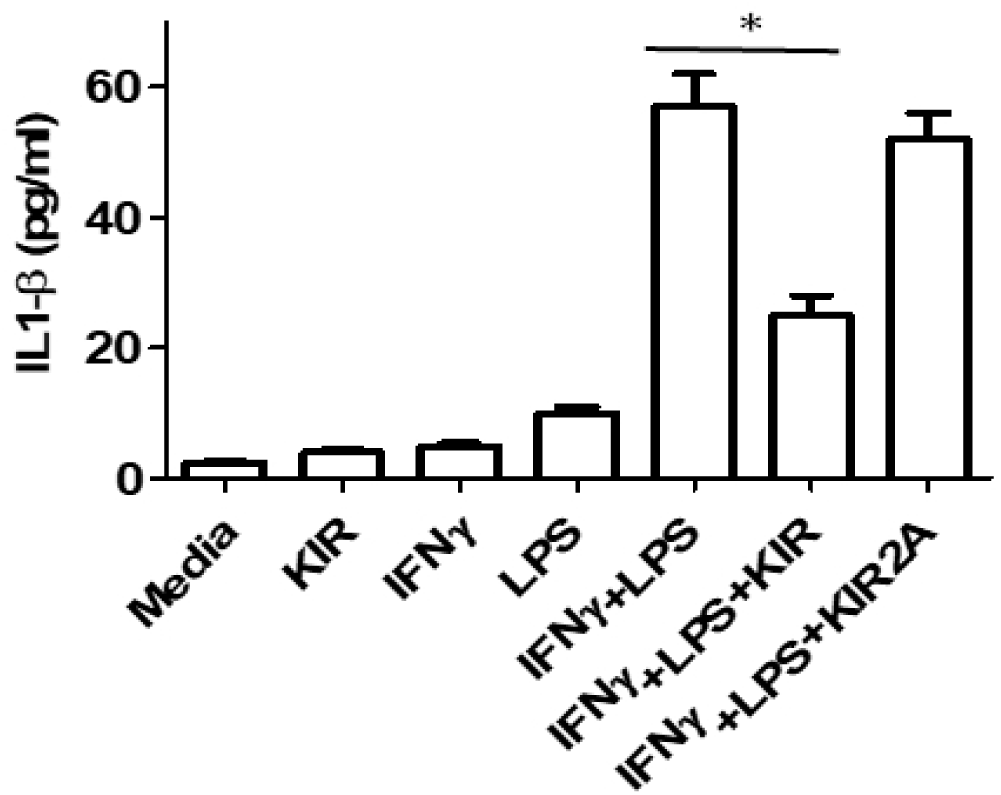
R9-SOCS1-KIR suppresses the IFNγ and LPS induced secretion of IL-1β from THP1 cells. THP1 cells were seeded in serum free media at a density of 3 × 10^5^ cells per well in 12 well plates and grown overnight. Cells were treated with R9-SOCS1-KIR (KIR) peptide, or its inactive control peptide R9-SOCS-KIR2A (KIR2A) at 20 μM for 3hr, followed by treatment with IFNγ (1 ng/ml) for 2 hr. LPS (1 μg/ml) was added to cells, after which they were incubated overnight. Cell supernatants were harvested and used for quantitation of IL-1β in an ELISA format. P = 0.01.

The NF-κB promoter is activated in response to inflammation caused by TNFα as well as by activation of toll like receptors TLR2 and TLR4. We therefore determined the effect of R9-SOCS1-KIR on the induction of NF-κB promoter by TNFα in ARPE-19 cells. Cells were pre-treated with R9-SOCS1-KIR or R9-SOCS1-KIR2A at 20 μM for 3 hr followed by treatment with TNFα at 50 ng/ml for 4 hr. Cells were then co-transfected with a mixture of plasmids containing NF-κB promoter linked to firefly luciferase and a thymidine kinase promoter driven Renilla luciferase as an internal control. Treatment with TNFα caused a 6.5-fold induction of NF-kB promoter, but induction was reduced by 50% in the presence of R9-SOCS1-KIR, while R9-SOCS1-KIR2A control peptide had negligible effect (**Figure 4**), indicating that R9-SOCS1-KIR was able to suppress the activation of NF-κB promoter.

**Figure 4.**
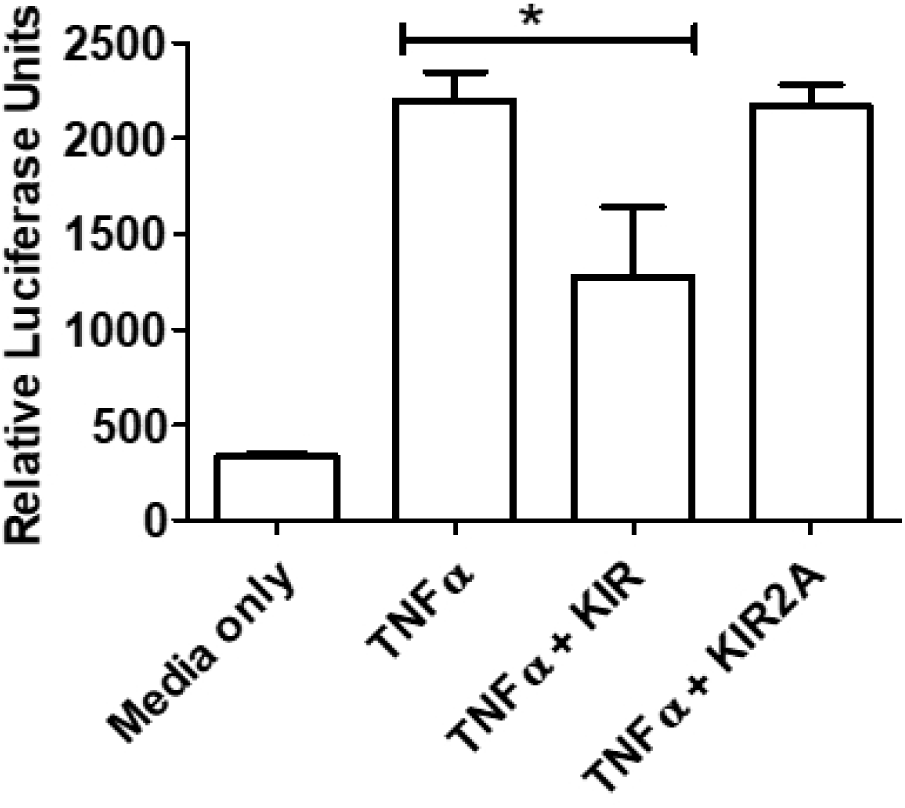
R9-SOCS1-KIR suppresses the induction of NF-kB promoter by TNFα. ARPE19 cells were seeded at about 80% confluency, grown overnight and changed to media without serum. They were treated with 20 μM of R9-SOCS1-KIR, or the control peptide R9-SOCS1-KIR2A for 3 hr, followed by addition of 10 ng/ml of TNFα for 4 hr. Cells were then co-transfected with a mixture of plasmids containing NF-kB promoter linked to firefly luciferase and a thymidine kinase promoter driving Renilla luciferase as an internal control for 24 hours. Cells lysates, in triplicate, were then used to measure relative luciferase activities. Average relative luciferase units are shown ± sd. *, p = 0.03.

### 3.3. Protection against disruption of barrier function of ARPE-19 cells by R9-SOCS1-KIR

A specialized group of proteins known as tight junction proteins provide a functional barrier lateral surface of RPE cells. We have used the inflammatory cytokines TNFα or IL-17A, which are known to cause disruption of barrier properties during uveitis to determine the protective effect of R9-SOCS1-KIR (Horai and Caspi, 2011). ARPE-19 cells were grown in low serum media for 4 weeks until they reached a cuboidal morphology similar to primary RPE cells. Cells were pre-treated with R9-SOCS1-KIR at 20 μM for 3 hr followed by TNFα (10 ng/ml), or IL-17A (50 ng/ml) for 48 hr. Cells were then stained with an antibody to zona occludin 1 (ZO-1) to observe the integrity of cell boundaries (**Figure 5**). Untreated cells or cells treated only with R9-SOCS1-KIR showed a continuous distribution of ZO-1 at the interface of neighboring cells. Treatment with TNFα or IL-17A caused a marked disruption of staining with ZO-1. This disruption was prevented when R9-SOCS1-KIR was simultaneously present in the cells.

**Figure 5.**
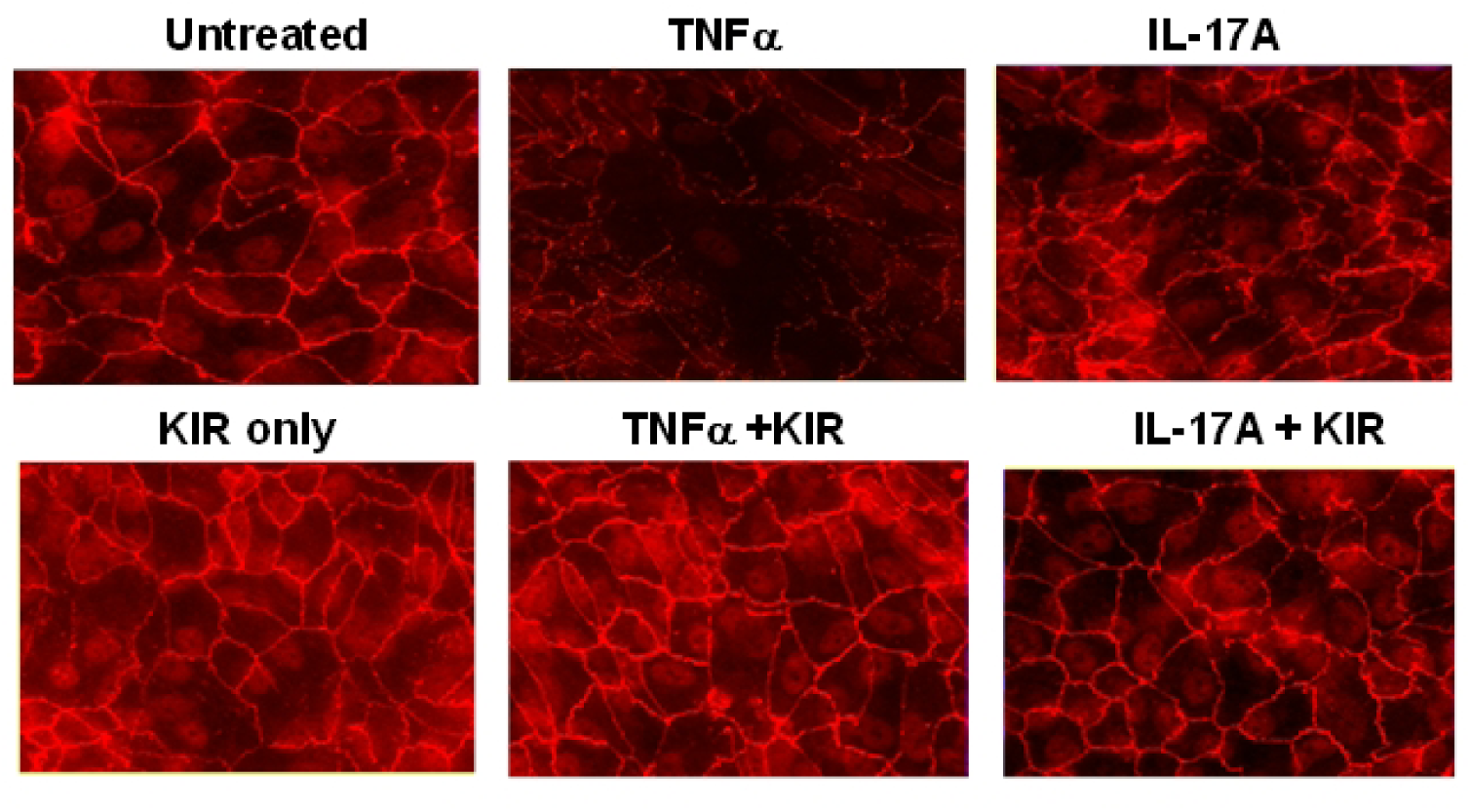
R9-SOCS1-KIR peptide protects ARPE-19 cell against the loss of tight junctions caused by TNFα or IL-17A. ARPE-19 cells grown in low serum media in 8 well slides for four weeks until they had differentiated into cuboidal shape. Cells were treated with R9-SOCS1-KIR, or its control inactive peptide, R9-SOCS1-KIR2A at 20 μM for 3 hr prior to addition of TNFα (10 ng/ml), or IL-17 (10 ng/ml) for 48 hrs. Untreated cells or those treated with TNFα or IL-17 were used alongside. Cells were stained with an antibody to ZO-1, followed by staining with a fluorescent conjugated secondary antibody, and imaged in a fluorescence microscope.

A consequence of loss of tight junction proteins is a reduction in the electrical resistance across the RPE monolayer. ARPE-19 cells were grown in transwell plates in low serum for 4 weeks. They were pre-treated with R9-SOCS1-KIR (20 μM) for 3 hr, followed by treatment with TNFα (10 ng/ml), or IL-17A (50 ng/ml) for 48 hr. Transepithelial electrical resistance (TEER) in cells was measured using a voltohmmeter (**Figure 6**). TEER observed in untreated cells was reduced by two-thirds by treatment with TNFα and by one-third by treatment with IL-17A. When R9-SOCS1-KIR was simultaneously present, the loss in TEER by TNFα or IL-17A was prevented, which is consistent with the preservation of tight junction protein observed with ZO-1 staining described above (**Figure 5**).

**Figure 6.**
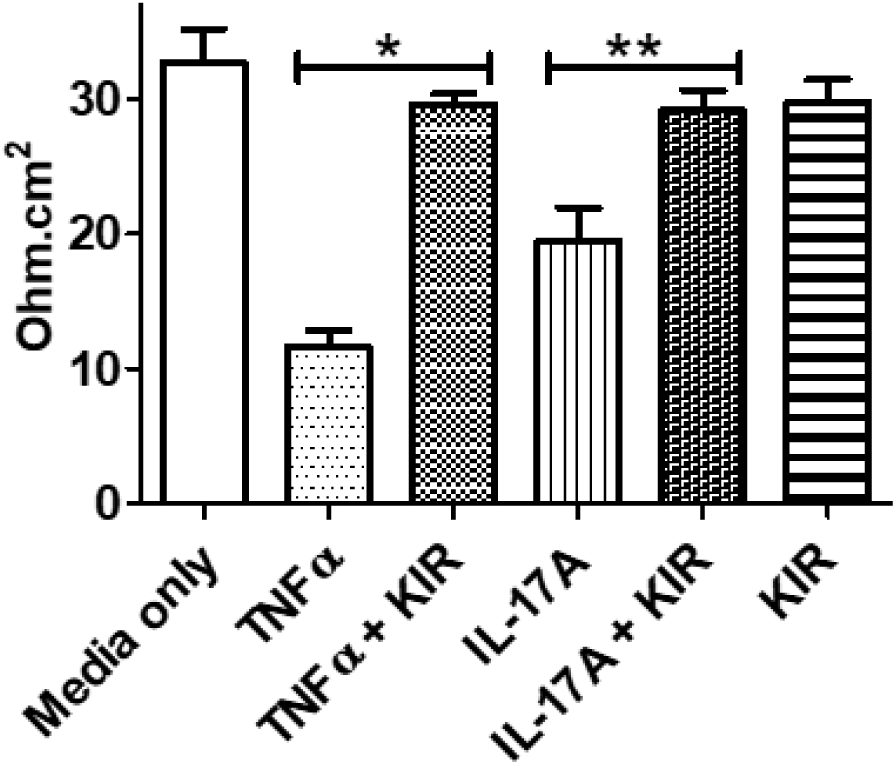
R9-SOCS1-KIR protects ARPE-19 cells against the loss of transepithelial electrical resistance (TEER) caused by inflammatory agents, TNFα and IL-17A. ARPE-19 cells grown in transwell plates (24 well) in low serum medium for 14 days were treated with R9-SOCS1-KIR (20 μM) for 3 hrs as indicated, followed by treatment with TNFα (10 ng/ml), or IL-17A (50 ng/ml) for 48 hr. Untreated cells or those treated with TNFα (10 ng/ml for 48 hr), or IL-17A (50 ng/ml for 48 hr) were used alongside. TEER in individual wells, in triplicate, was measured using EVOM2 voltohmmeter. *, p = 0.005; **, p = 0.009.

### 3.4. Topical administration of R9-SOCS1-KIR protects against damage from experimental autoimmune uveitis

B10.RIII mice (female, n=8) were immunized with a peptide derived from human inter photoreceptor binding protein IRBP (160-180) emulsified in complete Freund’s adjuvant. Pre-treatment with R9-SOCS1-KIR peptide (15 ug in 2 μl in each eye) or PBS (2 μl in each eye) began on day −2 and was performed daily for three weeks post immunization. Beginning two weeks after immunization, eyes were examined weekly by spectral domain optical coherence tomography (SD-OCT) and digital fundoscopy. A representative image by OCT is shown in **Figure 7, left panels**. While there were many infiltrating cells in the optical axis, vitreous as well as in retina of PBS treated eyes, there were significantly fewer cells in eyes treated with R9-SOCS1-KIR as eye drops. On day 21 after immunization, the average number of inflammatory cells in B-scans from three areas of the posterior chamber in PBS treated mice was 75 ± 13, while those scans of equivalent areas in R9-SOCS1-KIR treated mice was 3.5 ± 2 (p= 0.01, n=8). Retinal buckling and detachment were seen in PBS treated eyes and not in R9-SOCS1-KIR treated eyes. Similarly, a representative example of fundoscopy (**Figure 7, right panels**) shows the engorged blood vessels, deposits and hemorrhages in PBS treated eye, while the topical treatment with R9-SOCS1-KIR shows only a swollen optic nerve as the mild damage. A clinical score, calculated as follows was used to follow the disease: 0, no disease; 0.5, few infiltrating cells; 1, Many infiltrating cells; 2, 1 + engorged blood vessels; 2.5, 2 + hemorrhage, deposits; 3, 2.5 + retinal edema, detachment. **Figure 8** shows the clinical scores indicating the protective role of SOCS1-KIR-R9 versus PBS as eye drops in B10.RIII mice (n=8). Although the treatment with SOCS1-KIR-R9 was stopped at week 3, there was no further damage in these mice in the 2 weeks after the treatment was stopped.

**Figure 7.**
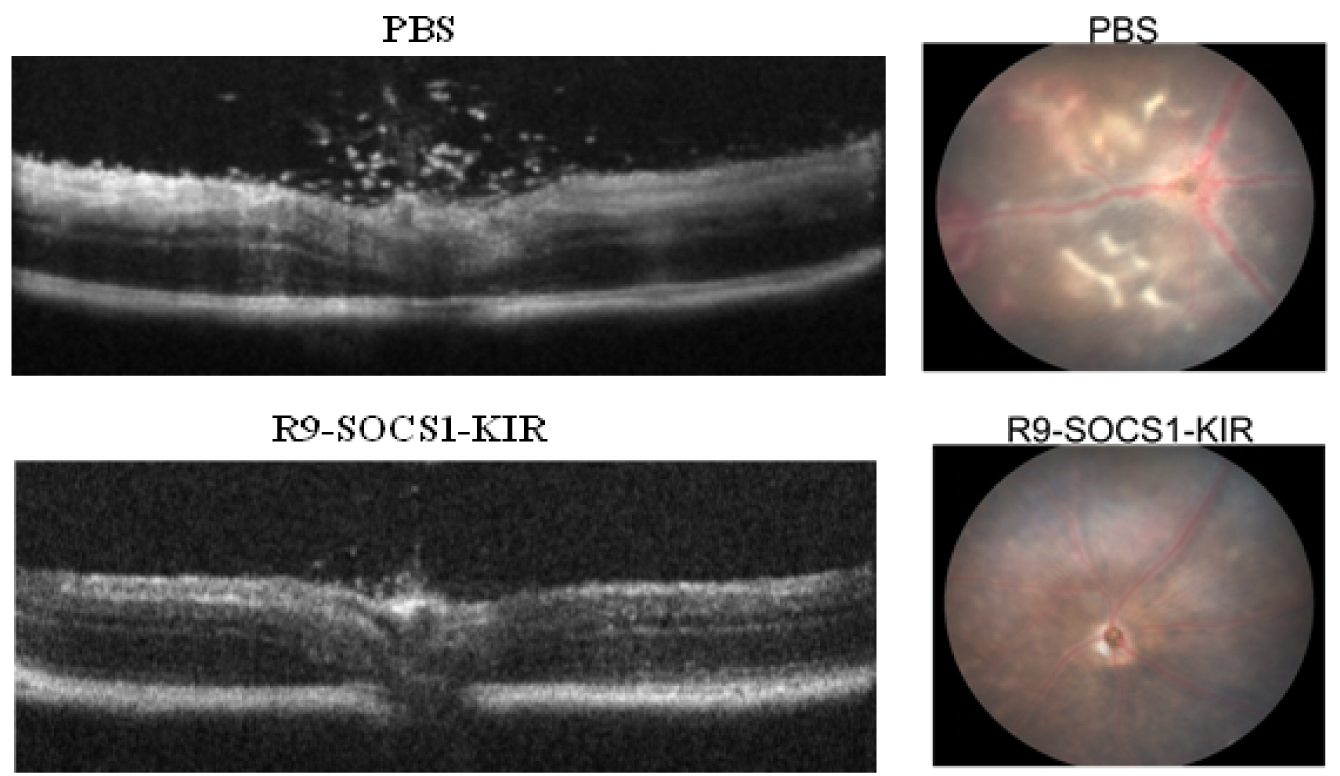
Protection against autoimmune uveitis in mice by topical administration of SOCS1-KIR-R9. B10.RIII mice (female, n=8) were administered sterile PBS or SOCS1-KIR-R9 (15 μg in 2 ul in each eye) on days −2, −1 and day 0. On day 0, mice were immunized with IRBP. Mice were treated daily with sterile PBS or R9-SOCS1-KIR (15 μg in 2 μl in each eye). A representative example of optical coherence tomography of eyes three weeks after the treatment is shown in the left panels. A decrease in infiltrating lymphocytes was noted in SOCS1-KIR-R9 treated mice (bottom), as opposed to the PBS treated mice (top). A representative example of fundoscopy of eyes on day 21, treated with sterile PBS (left), or R9-SOCS1-KIR for three weeks (right) is shown in the panels to the right. Note the engorged blood vessels, hemorrhage and deposits seen in PBS treatment were not seen upon R9-SOCS1-KIR treatment.

**Figure 8.**
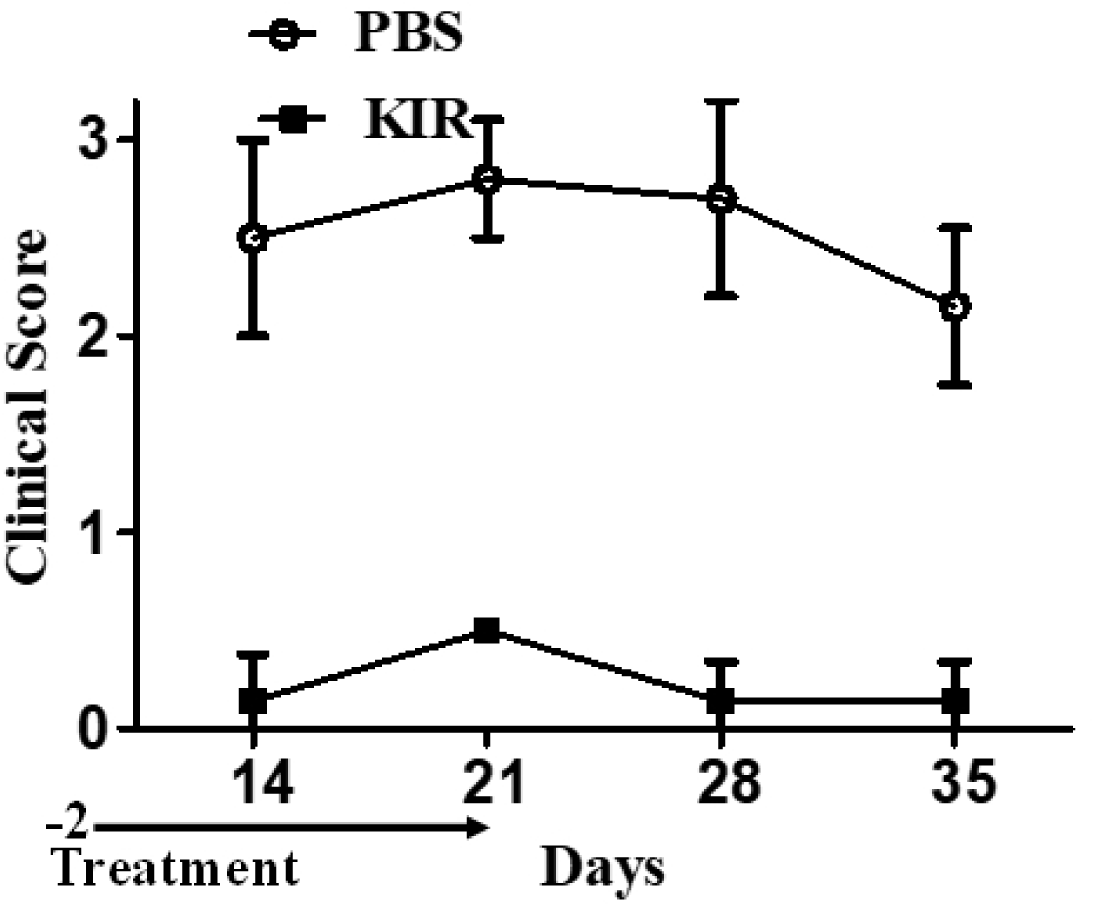
R9-SOCS1-KIR prevents the ocular damage of autoimmune uveitis. B10. RIII mice (female, n=8) were immunized with IRBP on day 0. Pre-treatment with PBS (open circles, 2 μl per eye), or R9-SOCS1-KIR (filled squares, 15 μg in 2 μl, per eye) as eye drops began on day −2 and continued until day 21. Examination of eyes by fundoscopy and optical coherence tomography was used to develop a clinical score as follows. 0, no disease; 0.5, few infiltrating cells; 1, Many infiltrating cells; 2, 1 + engorged blood vessels; 2.5, 2 + hemorrhage, deposits; 3, 2.5 + retinal edema, detachment.

To test if the presence of inflammatory cells and deposits noted above led to a decline in retinal function, we carried out electroretinography in dark-adapted mice before immunization and four weeks after immunization and the topical treatment with PBS or R9-SOCS1-KIR (**Figure 9**). Mice that were treated with PBS showed 60-70% decrease in amplitude across all intensities in a- and b- waves in both scotopic (dark-adapted) and photopic (light-adapted) electroretinograms, In contrast, mice that were treated with R9-SOCS1-KIR for 3 weeks showed only a 10% decrease in scotopic ERGs, while the photopic ERG amplitudes that reflect the function of the cones were fully preserved.

**Figure 9.**
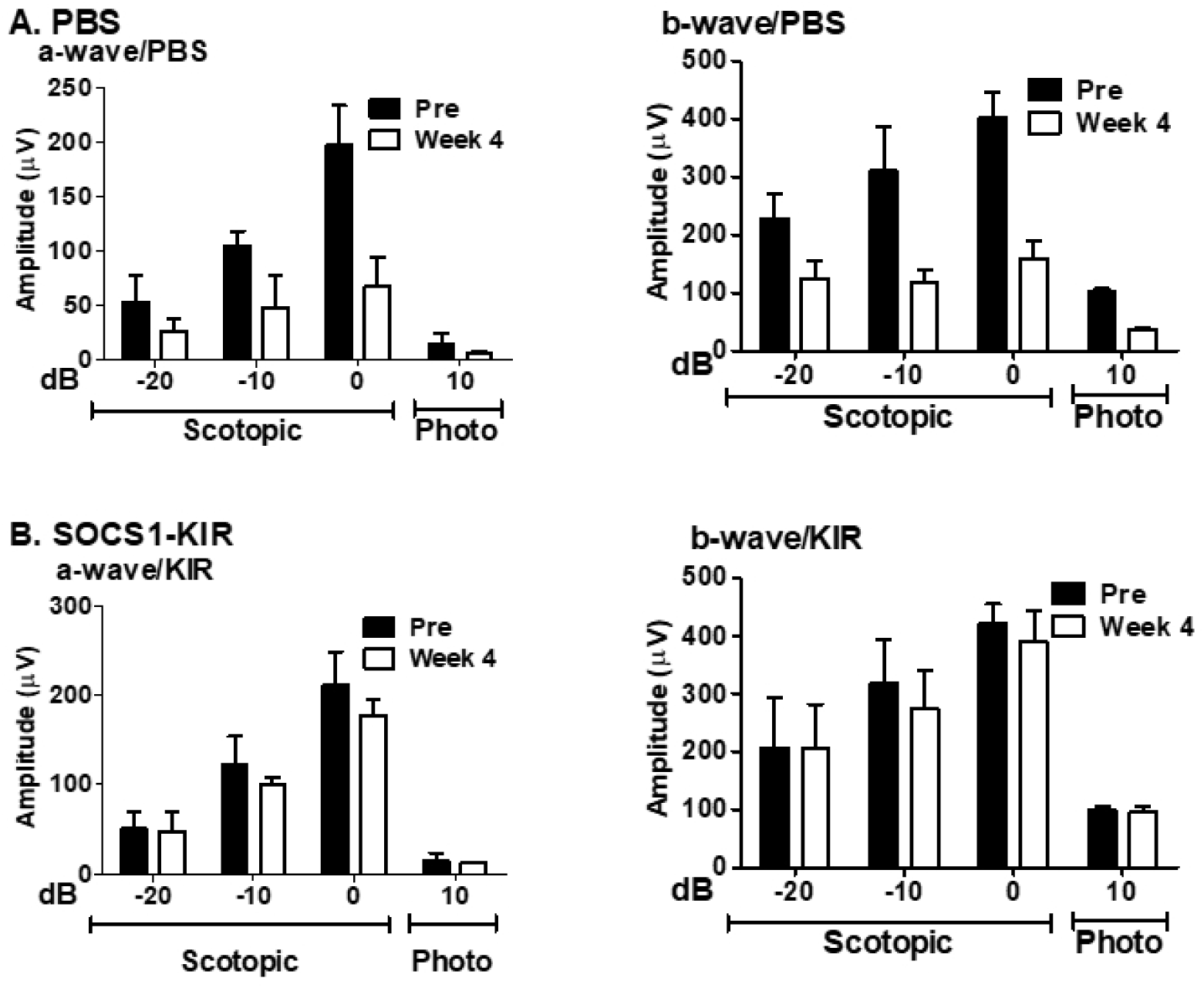
Topical treatment with R9-SOSC1-KIR protects against loss of a-wave and b-wave amplitudes. Dark-adapted and light-adapted electroretinogram responses were measured in B10.RIII mice before immunization (Pre), and 4 weeks after immunization with IRBP peptide and topical administration of PBS (**A**, top lanes), or SOCS1-KIR (**B**, bottom lanes, abbrev: KIR). Average a-wave amplitudes and b-wave amplitudes (scotopic at −20, −10 and 0 dB, and photopic, 10 dB) are shown. The same cohort of mice were used for all data points.

## 4. Discussion

SOCS1 is an important negative regulator of JAK/STAT as well as NF-KB signaling. SOCS-1 carries a SOCS box domain that directs the complex consisting of the ligand, receptor, JAK kinase and associated STAT to the proteasome for degradation. We have shown previously that lipo-SOCS1-KIR can by itself recapitulate properties of intact SOCS1 and protect mice against experimental autoimmune encephalomyelitis (EAE) (Jager et al., 2011). Because it lacks the SOCS box, SOCS1-KIR does not direct the ligand/receptor complex to the proteasome for degradation and thus has a longer half-life than intact SOCS1. In contrast to the lipo-SOCS1-KIR used previously, in this study we synthesized and tested a polyarginine (R9) attached SOCS1-KIR (R9-SOCS1-KIR), which offers several advantages: It is water soluble and thus the need to use dimethylsulfoxide (DMSO) for solubilization of lipo-SOCS1-KIR is eliminated. Next, absence of a lipid moiety makes the molecule less immunogenic. Lipo-SOCS1-KIR originally reported by Jager et al. (Jager et al., 2011) has been used by others in C57BL/6 mice (He et al., 2015) and Lewis rats (He et al., 2016) in protection against EAU. The R9-attached peptide is thus an improvement over the lipo-mimetic (Jager et al., 2011) for use of KIR peptide. The utility of SOCS1-KIR-like peptides in the treatment of other autoimmune diseases has also been reported in diabetes (Recio et al., 2014), and in psoriasis (Kubo, 2013).

Cell penetrating peptides are an important tool for the delivery of therapeutic compounds (Copolovici et al., 2014; Kristensen et al., 2016). Blood-brain crossing cell penetrating peptide for vascular endothelial growth inhibitor (VEGI) have also been described (Gao et al., 2015). Eye drop formulations of pigment epithelium derived factor (PEDF) peptide (Liu et al., 2012; Vigneswara et al., 2015), and endostatin (Zhang et al., 2015) have also been tested. The topical delivery of R9-SOCS1-KIR we describe here seems to be applicable for other peptides as well.

While steroid therapy is one of the current lines of treatment of autoimmune uveitis and is helpful for anterior uveitis, it is not effective for panuveitis. In contrast to the intravitreal injections required and the undesirable side effects associated with steroid therapy, topical administration of R9-SOCS1-KIR is an straightforward alternative for panuveitis. To confirm its utility for human disease, future experiments should demonstrate that R9-SOCS-KIR can access the anterior chamber and the posterior chamber in a larger animal model such as the rabbit (Edmond et al., 2016).

SOCS1-KIR binds to the activation loop of tyrosine kinases JAK2, TYK2, and the adaptor protein MAL (Jager et al., 2011; Waiboci et al., 2007). SOCS1-KIR binding to JAK2 and TYK2 limits the extent and duration of immune response generated by the cytokines from T helper 1 cells (IFNγ, e.g.), and from T helper 17 cells (IL17A, e.g.) cells. Simultaneous inhibition of both Th1 and Th17 responses in uveitis is critical, since inhibition of one of these cytokines exacerbates the effect of the other (Luger et al., 2008). Binding to the adaptor protein MAL, on the other hand, limits the inflammatory signaling initiated from toll like receptors 2 and 4 (TLR2 and TLR4) by interfering with MyD88 signaling, leading up to suppression of NF-kB pathway. SOCS1 binding to the p65 subunit of NF-kB complex downregulates the signaling NF-kB promoter (Strebovsky et al., 2011). Furthermore, the binding of SOCS1-KIR to Rac1-GTP suppresses the generation of reactive oxygen species (ROS), and thereby suppresses the activation of inflammasome NLRP3 (Inoue and Shinohara, 2013). TNFα, which is another inflammatory cytokine that is elevated during uveitis acts through NF-kB promoter, the activity of which is suppressed by SOCS1-KIR. SOCS1-KIR is thus capable of suppressing a multitude of inflammatory responses.

Binding of R9-SOCS1-KIR to phosphotyrosine residues of tyrosine kinases JAK2 and TYK2 leads to the downregulation of the activation of transcription factors STAT1 and STAT3, respectively (**Fig 1**). Suppression of STAT1 and STAT3 leads to downregulation of the effects of the inflammatory cytokines IL-6 and IL-22 (a cytokine produced by Th17 cells), both of which are elevated in uveitis and related autoimmune disorders (Kim et al., 2016; Sainz-de-la-Maza et al., 2016). Uveitis patients were shown to have elevated levels of IL-22 (Takeuchi et al., 2017). Since IL-22 utilizes JAK2 and TYK2 for its signaling, the effect of this cytokine is likely to be dampened in the presence of R9-SOCS1-KIR. Importance of STAT3 is further supported by the observation that genetic deletion of STAT3 in CD4 cells leads to a resistance to autoimmune disorders (Harris et al., 2007; Liu et al., 2008). Binding of SOCS1-KIR to the adaptor protein MAL dysregulates signaling from MyD88 and NF-kB. It is worth noting here that genetic deletion of either MyD88 or NF-kB leads to a resistance to autoimmune disorder (Mc Guire et al., 2013; van Loo et al., 2006).

There is cross talk between SOCS1 and regulatory T cells (Tregs), such that the protective Tregs are maintained in the presence of SOCS1 (Ahmed et al., 2015; Chang et al., 2012). SOCS1 and SOCS1-KIR prevent the loss of Foxp3, which is required for the formation of Tregs. Some of the Tregs can be converted into Th1- or Th-17-like cells, which is prevented by SOCS1 and SOCS1-KIR. For these reasons, SOCS1 has been regarded as the ‘guardian’ of Tregs (Yoshimura et al., 2012). In addition, SOCS1 has been shown to enhance M2 macrophages (neuroprotective), while simultaneously decreasing M1 subtype (inflammatory) (Whyte et al., 2011; Wilson, 2014).

Mouse models of genetic deletion of specific signaling molecules provide further evidence of the usefulness of R9-SOCS1-KIR described above. Mouse models of deletion of MyD88 (Su et al., 2005), the adaptor partner of MAL binding repressed by SOCS1-KIR are resistant to EAU. Similarly, we showed that R9-SOCS1-KIR inhibited STAT3 signaling. Mice deleted in STAT3 in CD4 cells are resistant to EAU (Liu et al., 2008). Loss of NF-kB in CNS ameliorates autoimmune encephalomyelitis in mice (Mc Guire et al., 2013; van Loo et al., 2006).

RPE plays many important roles including the maintenance of blood-ocular barrier, providing nourishment to photoreceptors and removal of waste products. Although the beneficial effects of R9-SOCS1-KIR shown above were carried out in ARPE-19 cells, the immune suppressive effects of this therapeutic peptide should play out in all cells that have an active JAK/STAT and/or NF-kB signaling system, which includes the activated microglia and the invading macrophages and T cells during the course of uveitis. The anti-inflammatory effects and preservation of barrier functions of R9-SOCS1-KIR noted in ARPE-19 cells may have contributed to the protection of retinal structure and function in B10.RIII mice immunized with IRBP peptide. Inflammatory cells in the optical axis, vitreous and retina were noted on OCT analysis. Deposits, engorged blood vessels and hemorrhage were observed in fundus examination of uveitic mice. A decrease in retinal function as documented by a significant decrease in scotopic and photopic a- and b-wave of ERG recordings at all light intensities were observed in uveitic mice treated only with vehicle. Topical treatment with R9-SOCS1-KIR protected mice against the structural and functional damage associated with IRBP induced uveitis, as evidenced by the milder disease indications and clinical scores. R9-SOCS1-KIR may thus be a potential treatment for uveitis and other inflammatory ocular diseases, such as macular degeneration.

## 5. Conclusions

R9-SOCS1-KIR was effective in preventing ocular damage in a mouse model of experimental autoimmune uveitis. Trafficking of the infiltrating cells, hemorrhage and other retinal damages seen in control mice were not observed upon topical administration of R9-SOCS1-KIR. Suppression of IFNγ and IL-17A signaling, downregulation of chemokines that recruit inflammatory cells as well as preservation of barrier properties of RPE noted in ARPE-19 cells may have contributed to the protection seen in B10.RIII mice. Given the many overlapping features of infectious and autoimmune uveitis, R9-SOCS1-KIR is a potential candidate for the treatment of infectious uveitis as well. Individuals with HLA-29 haplotype are prone to developing certain forms of uveitis. It is conceivable that a prophylactic treatment with R9-SOCS1-KIR in these patients as they age will slow the progression of their disease. R9-SOCS1-KIR may also be a potential therapeutic for autoimmune keratitis and age related macular degeneration.

## Acknowledgments

This work was supported by Shaler Richardson Professorship endowment to ASL and by the NEI core grant to the University of Florida (P30 EY02172). We thank Dr. Eduardo Candelario-Jalil for allowing us to use his voltohmmeter to measure transepithelial electrical resistance.

